# Heat stress reveals high molecular mass proteasomes in *Arabidopsis thaliana* suspension cells cultures

**DOI:** 10.1101/501031

**Authors:** Daniel Aristizábal, Viridiana Rivas, Gladys Cassab, Fernando Lledías

**Affiliations:** Departamento de Biología Molecular de Plantas, Instituto de Biotecnología, Universidad Nacional Autónoma de México, Av. Universidad 2001, Col. Chamilpa, Cuernavaca, Mor., 62250, México

**Keywords:** *Arabidopsis* thaliana suspension cell culture, Proteasome, Heat stress, Blue-Native gel electrophoresis

## Abstract

Because of their sessile nature, plants have evolved complex and robust mechanisms to respond to adverse environments. Stress conditions trigger an increase in protein turnover and degradation. Proteasomes are essential to the cell for removing, in a highly regulated manner, partially denatured or oxidized proteins thus minimizing their cytotoxicity. We observed that suspension cells of *Arabidopsis thaliana* treated with high temperature (37 °C) directed the assembly of high molecular mass proteasomes. The removal of a 75% of the original ubiquitin conjugates and the maintenance of protein carbonyls at basal levels correlated with a specific proteasome profiles. The profiles obtained by the separation of different proteasomes populations by Blue-Native Polyacrylamide Gel Electrophoresis and western blot analysis suggest that synthesis, assembly, and heavy ubiquitination of 20S (CP) subunits are promoted by heat stress.

## 1. Introduction

Plants undergo various stressful environments across their lifetimes, but their sessile nature means they cannot escape from unfavorable conditions. Plants have developed unique strategies for stress mitigation or adaptation to their surroundings. Since stress environments activate an increase in protein turnover and degradation, one strategy to cope with it is the selective protein breakdown mediated by the proteasome in the nucleus, cytosol and endoplasmic, reticulum which decreased their toxicity (Smalle and Vierstra, 2004), (Thompson and Vierstra, 2005). From *in vitro* studies, it was shown that the 20S proteasome actively recognizes and degrades oxidized proteins, in contrast to the 26S proteasome, which is not very effective even in the presence of ATP and the ubiquitination system (Shang and Taylor, 1995), (Obin et al.,1998). This may be explained by the fact that a mild oxidative stress rapidly inactivates both the ubiquitin-activating-conjugating system and 26S proteasome activity in intact cells but does not affect 20S proteasome activity (Davies, 2001).

Proteasomes are protein degradative complexes involved in all processes of the living cell such as cell division, stress response, transcription, DNA repair, and signal transduction, among others (Glickman and Ciechanover, 2002; Hershko and Ciechanover, 1998). In plants, proteasomes have been particularly involved in the differentiation of leaves, flowers, and xylem, in hormone response, as well as abiotic and biotic stress responses (Shibahara et al., 2002). Proteomic analysis of different organism reported that an estimated 80 to 90% of the cytosolic proteins are degraded via proteasomes (Glickman and Ciechanover, 2002). The minimal expression of a proteasome is the 20S core particle, or catalytic particle (CP) constituted by four stacked seven-membered rings of β _1-7_ (central) and α _1-7_ (distal) subunits in an arrangement α-β-β-α, for a total of 28 subunits. In this hollow-cylinder structure, the interior domains of the subunits β1, β2 and β5 define catalytic domains that have trypsin, chymotrypsin, and caspase-like activities respectively (Kish-Trier and Hill, 2013). Virtually any protein in direct contact with the proteasome catalytic sites could be degraded (Nussbaum et al., 1998). The first line of proteasome regulation, that limits the inappropriate and non-selective protein degradation, are the N-terminus regions of α subunits that constitute the proteasome gate (Groll et al., 2000). This basic 20S (CP) proteasome version is responsible for cell removal of oxidatively modified proteins (Davies, 2001). The function and structure of the 20S (CP) is highly conserved through eukarya (Tanaka et al., 1988; Fort et al., 2015). 20S α subunits are also the binding sites for different regulatory complexes. The 26S proteasome for example, is formed by the basic 20S (CP) flanked by one or two 19S regulatory complexes (19S-20S or 19S-20S-19S) docked on the distal-most surfaces of the α rings. For *Saccharomyces cerevisiae, Schizosaccharomyces pombe* and human cells, the architecture of the 19S regulatory particle has been reported at sub-nanometer level and the functions of most of the subunits has been assigned (Beck et al., 2012; Schweitze et al., 2016; Sakata et al., 2012; Bohn et al., 2010). As for the independent 20S (CP), the structure and function of the 19S regulatory complex is conserved trough evolution (Fort et al., 2015). The 19S complex is in charge of recognizing ubiquitinated proteins, denature them by the action of ATP-dependent “unfoldases”, opens the 20S (CP) gate by rearrangement of the α subunits, and translocate the protein substrate to the catalytic sites for its degradation (Peth et al., 2010). The 26S is the specialized proteasome version involved-along with a ubiquitin activating (E1), the ubiquitin conjugating (E2) and the ubiquitin ligases (E3) enzymes-in the Ubiquitin-Proteasome-System (UPS) (Hershko and Ciechanover, 1998). Approximately 5% of the *A. thaliana* proteome is dedicated to this system or its regulation (Manzano et al., 2008). A different proteasome regulatory particle PA28 (11S, REG), is a trimeric complex that degrades carbonylated proteins in an ATP independent manner (Hernebring et al., 2013). PA200 (in mammals) and the homologous Blm10 (in yeast), are alternative regulatory complexes which had been involved in the degradation of very specific non-ubiquitinated protein substrates (López et al., 2011; Blickwedehl et al., 2012). Ecm29 (in a human cell line) has been proposed to act as a structural stabilizing agent for the 26S proteasome, especially when the 20S (CP) maturation was impaired (Lehmann et al., 2010). Information on different proteasome regulators published so far has shown that the binding of the regulatory complexes has a deep influence in the selection of substrates, in the catalytic function of the 20S (CP) and in the peptides produced. In this scenario, the presence of “supra-20S” complexes allow the cell to respond to very specific metabolic stages. The active building of proteasomal complexes has a counterpart, the 26S proteasomes in yeast were disarmed into independent 19S and 20S (CP) particles during the stationary phase and this phenomenon correlated with cell viability. When ATP was available by medium refreshing, the 26S proteasomes were reconstituted (Bajorek et al., 2003). Dissociation and reassociation of the 26S proteasome have been reported during the adaptation of a human cell line to oxidative stress (Grune et al., 2011). Reversible 26S disassembly has been reported upon mitochondrial stress in yeast (Livnat-Levanon et al., 2014). Considering the assembly/disassembly capacities of the proteasomes and the repertoire of regulatory complexes, any cell in a specific environmental context, have the possibilities to “direct” the versions of the proteasome that respond better to the catalytic needs of the moment, and the proteasome versions selected would have an influence on cell fate. This information indicates that a very dynamic process of assembly, selection of regulatory particles, and disassembly of proteasomes are continuously taking place in cells in close relationship with the environment. Great efforts have been made to characterize biochemically, genetically and by mixed “omics” techniques, the subunit composition and function of the proteasomes in plants. In *A. thaliana* particularly, their 20S (CP) and 26S proteasomes have been isolated and characterized (Polge et al., 2009; Yang et al., 2004; Book et al., 2010). Unique genes code for the 20S (CP) and the 19S subunits in yeast and mammalian cells, in *A. thaliana* instead, both complexes are encoded by gene pairs whose sequence differences, suggest protein products with altered functions (Yang et al., 2004), so potentially different 20S (CP), 26S particles and probably other proteasomes entities assembled with paralogous pairs coexist. By proteomic approach using an epitope-tagged 26S proteasomes as bait, more than 40 proteins interact with this complex. This in-depth mass spectrometric analysis strongly suggests the existence of a diverse array of proteasomes. Despite all the information on *A. thaliana* proteasomes, studies on the dynamics of the different proteasome versions in plants as has been reported for yeast (Bajorek et al., 2003; Livnat-Levanon et al., 2014) or mammalian cells (Grune et al., 2011; 34. Shibatani et al., 2006) are limited.

Blue Native Polyacrylamide Gel Electrophoresis (BN/PAGE) is a technique that allows the separation of native protein complexes based on their molecular mass differences (Wittig et al., 2006). BN/PAGE has been used for the complexomics analysis of different models (Wittig et al., 2006; Lasserre et al., 2006; Hashemi et al., 2016). This technique has been employed to successfully separate the different versions of proteasomes in whole cell lysates of a human embryonic cell line (HEK293) (Camacho-Carvajal et al., 2004) and to study the proteasome dynamics of rabbit reticulocytes (Shibatani et al., 2006). In the latter report, six native proteasome populations (20S, 20S-PA28, PA28-20S-PA28, 19S-20S-PA28, 19S-20S and 19S-20S-19S) were identified. By γ-interferon stimulation or the chemical inhibition of the proteasome, an active interchange of proteasome regulatory “caps” was evidenced (Shibatani et al., 2006). The applicability of BN/PAGE in combination with label-free protein quantification and protein correlation profiling was employed to investigate the 20S proteasome from *Plasmodium falciparum,* even in the background of the whole protein extract (Sessler et al., 2012). BN/PAGE was used for monitoring changes in the quantity and subunit composition of the 20S (CP) when the α3 subunit was deleted in yeast (Couttas et al., 2011).

In the present work, we adapted some of the above-mentioned protocols of proteasome isolation and analysis by BN/PAGE to establish whether in *A. thaliana* cells in suspension culture different proteasome versions coexist and if under drastic changes in the culture conditions like heat stress, the basal proteasome populations were altered. Our results showed that cells in suspension culture subjected to an increment of temperature (from 25 to 37^°^ C) experienced an enrichment of high molecular mass proteasomes that in turn have an impact on the cell content of oxidized and ubiquitinated proteins.

## 2. Methods

### 2.1 Cell culture and heat stress treatment

Suspension cell cultures were generated from hypocotyls dissected from *A. thaliana* seedlings and were kindly provided by P. Guzmán and L. Aguilar (CINVESTAV, Irapuato). Cells were maintained by weekly transfer in MS medium (Murashige and Skoog, 1962) containing basal salt mixture, 3% sucrose and supplemented with 50 μg /L kinetin, 75 μg/L 2,4-diclorofenoxiacetic acid (2-4D) and 1X Gamborg’s vitamin solution, pH 5.7. Cultures were incubated at 25 °C and 100 rpm under long day conditions of 16 h light/ 8 h dark, and 80 µM photons m^-2^ s^-1^. For heat stress treatment, a one-week “mother culture” of exponentially growing cells was diluted (1:10) with fresh MS medium (plus supplements) and divided into independent 250 mL flasks containing 50 mL liquid medium. Cell cultures were incubated at 37 °C at 100 rpm for 0.5, 1, 2 and 3 h (illuminated), then individual cultures were immediately filtered using regular paper towels to discard liquid medium. Cell packages (approximately 10 mL) were recovered with a spatula and immediately frozen in liquid nitrogen. Samples were kept at −70 °C until their processing for proteasome isolation.

### 2.2 Proteasomes isolation

Total cell lysate was obtained by adding to each frozen cell package, 25 mL of extraction buffer (Tris-HCl pH 7.5, 1 mM dithiothreitol, 2 mM adenosine triphosphate, 0.25M sucrose, 1 mM MgCl2, 1% polyvinylpolypyrrolidone, Complete EDTA-free [Roche, used as recommended]), and “10 mL” of glass beads (4 mm). While thawing, cells were disrupted by five vortex cycles (5 min vortexing followed by 5 min incubation in ice). At this stage, aliquots from different cultures were taken for the determination of the total content of ubiquitin conjugates and protein carbonyls by western blot (see below). Total lysate was filtered on three layers of cheesecloth to retire glass beads and was centrifuged at 16 000 X g for 15 min at 4 °C. Pellet (P1, Fig. 1) was eliminated and supernatant (Sn1, Fig. 1) was centrifuged for 1h at 70 000 X g at 4 °C. Resultant pellet (P2, Fig. 1) was discarded and supernatant (Sn2, Fig. 1) was centrifuged again, this time at 350 000 X g for 3.5 h at 4 °C. The supernatant (Sn3, Fig. 1) was discarded, the pellet (P3, Fig.1) which contained the proteasomes enriched fraction, was resuspended in buffer A (HEPES buffer pH 7.8, 75 mM NaCl, 375 mM MgCl2, 40 mM DTT, glycerol 7.5% y 1.6 µM ATP). Aliquots were immediately separated by BN-PAGE or kept at −70 °C until their analysis.

**Fig. 1.**
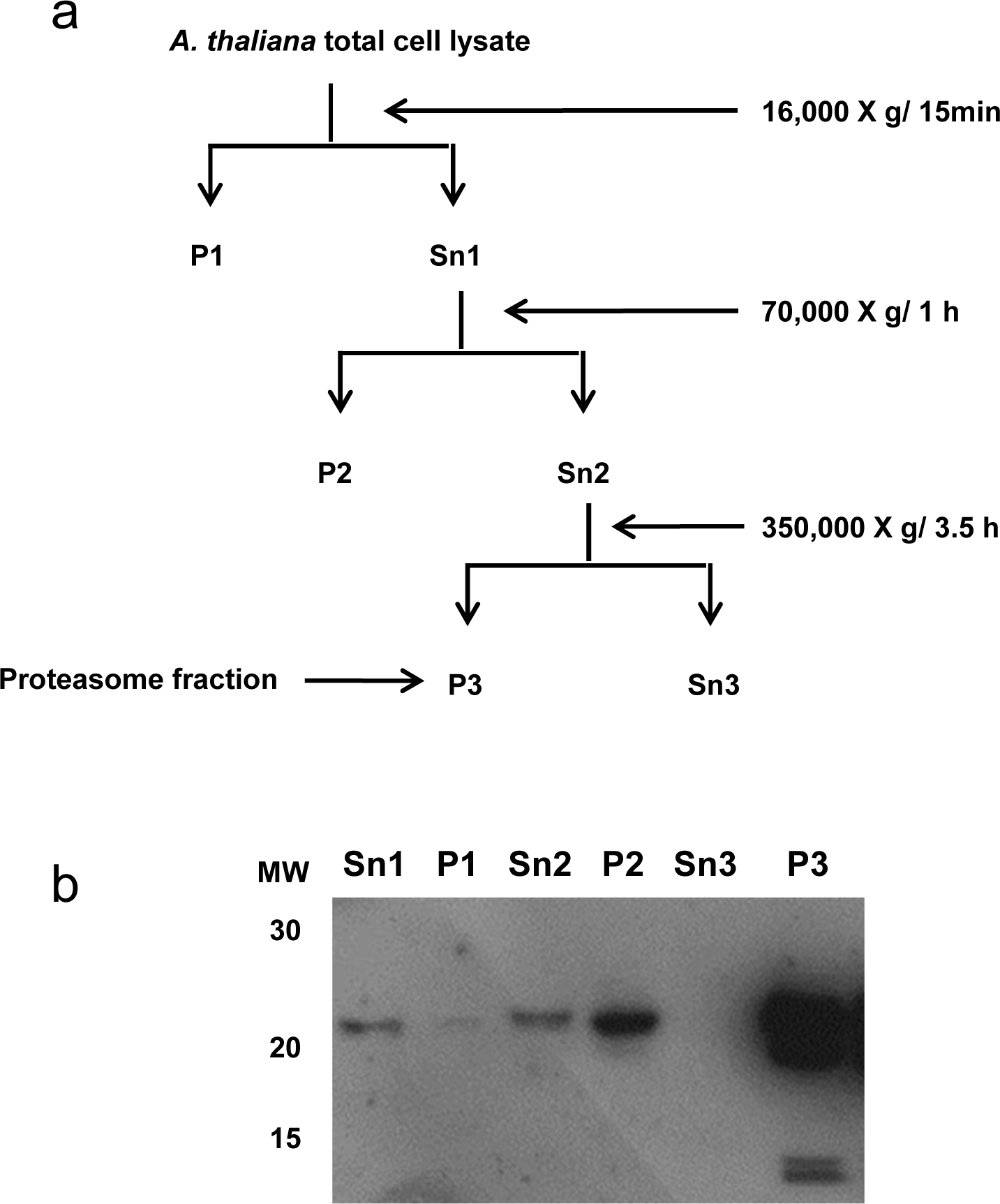
Proteasomes isolation scheme. A proteasome enriched fraction was obtained by differential centrifugation from total lysates of *A. thaliana* cell suspension cultures (a). Representative aliquots of all the pellets (P1 to P3) and supernatants (Sn1 to Sn3) were separated by SDS/PAGE and analyzed by western blot using an anti-20S antibody to determine the protocol efficiency (b). The proteasome enriched fractions (P3) resuspended in buffer A were directly loaded onto BN/PAGE gels to separate the proteasomes based on their molecular mass differences.

### 2.3 BN/PAGE

Resolution of proteasomes according to their molecular mass was achieved by BN/PAGE (Wittig et al., 2006), and optimized for proteasome analysis (Camacho-Carvajal et al., 2004; Shibatani et al., 2006) with some modifications. Proteasomes enriched fraction in buffer A was directly loaded onto an 8 × 6 cm BN/PAGE mini gels (5 to 10% acrylamide gradient [acrylamide:bis-acrylamide 32:1] in 50 mM BisTris/HCl, pH 7.0, 500 mM α-aminocaproic acid, and overlaid by 4% stacking gel in the same buffer). Electrophoresis was carried out at 5 °C according to the program: 50 V for 1h, 150 V for 16h and 500 V for 1h. Cathode buffer: 50 mM Tricine, 15 mM BisTris-HCl pH 7.0 and 0.02% Coomassie G-250 (Cat. 1442C-1, Research Organics, Inc.), anode buffer: 50 mM Bis-Tris-HCl pH 7.0 in a Mini-PROTEAN System (Bio-Rad). Proteins in analytical BN/PAGE were visualized with Coomassie Brilliant Blue (section 2.5) or by silver stain [40] (Blum et al., 1986). For preparative purposes, BN/PAGE gels were fractionated and electroeluted (next section). The molecular mass of the protein complexes was estimated by the method of Wittig et al., 2010, using the endogenous HSP 60, Rubisco (both proteins identified by mass spectrometry in the fraction 4 and 7 respectively), and the independent 20S (CP) native complexes as molecular mass markers.

### 2.4 Protein electroelution and concentration

After electrophoresis running completion, a glass plate of the gel “sandwich” was removed. As the cathode buffer contained Coomassie G-250, four protein bands were stained during electrophoresis (Fig. 2b) and used as markers to cut the gels into eight horizontal fractions (Fig. 2c). For the analysis of the disassembled proteasomes components by western blot, each fraction was individually divided into smaller fragments (∼ 2 × 2 mm) and heated at 95 °C for 10 min in the presence of 1 mL of 1X Laemmli sample buffer (Tris 50 mM pH 6.8, 2% SDS, 5% β-ME, 8% glycerol without bromophenol blue). After cooling, gel pieces and sample buffer were transferred to the sample traps of the electroelutor/concentrator system “Little Blue Tank” (ISCO, INC) containing 10 mL of Laemmli running buffer (25 mM Tris, 192 mM glycine, 0.1% SDS) at a 1:10 dilution. Laemmli running (1X) buffer was used in both electrode compartments. Protein electroelution was carried out at 3 W for 3 h at 5 °C. Samples recovered from the sample traps (∼ 200 μL) were precipitated by methanol/chloroform, air dried, resuspended and heated at 95 °C for 10 min in 1X Laemmli sample buffer for their SDS/PAGE and western blot analysis. A similar technique was followed to recover native proteasome complexes from BN/PAGE, except that gel fractions were not heated or incubated in the presence of chaotropic agents. For this purpose, electroelutor/concentrator sample traps contained 10 mL of running buffer (1:10) and 1X running buffer was used in electrode compartments (without SDS). Samples recovered were immediately loaded or preserved at −70 °C after glycerol addition (20% final) for additional BN/PAGE analysis.

**Fig. 2.**
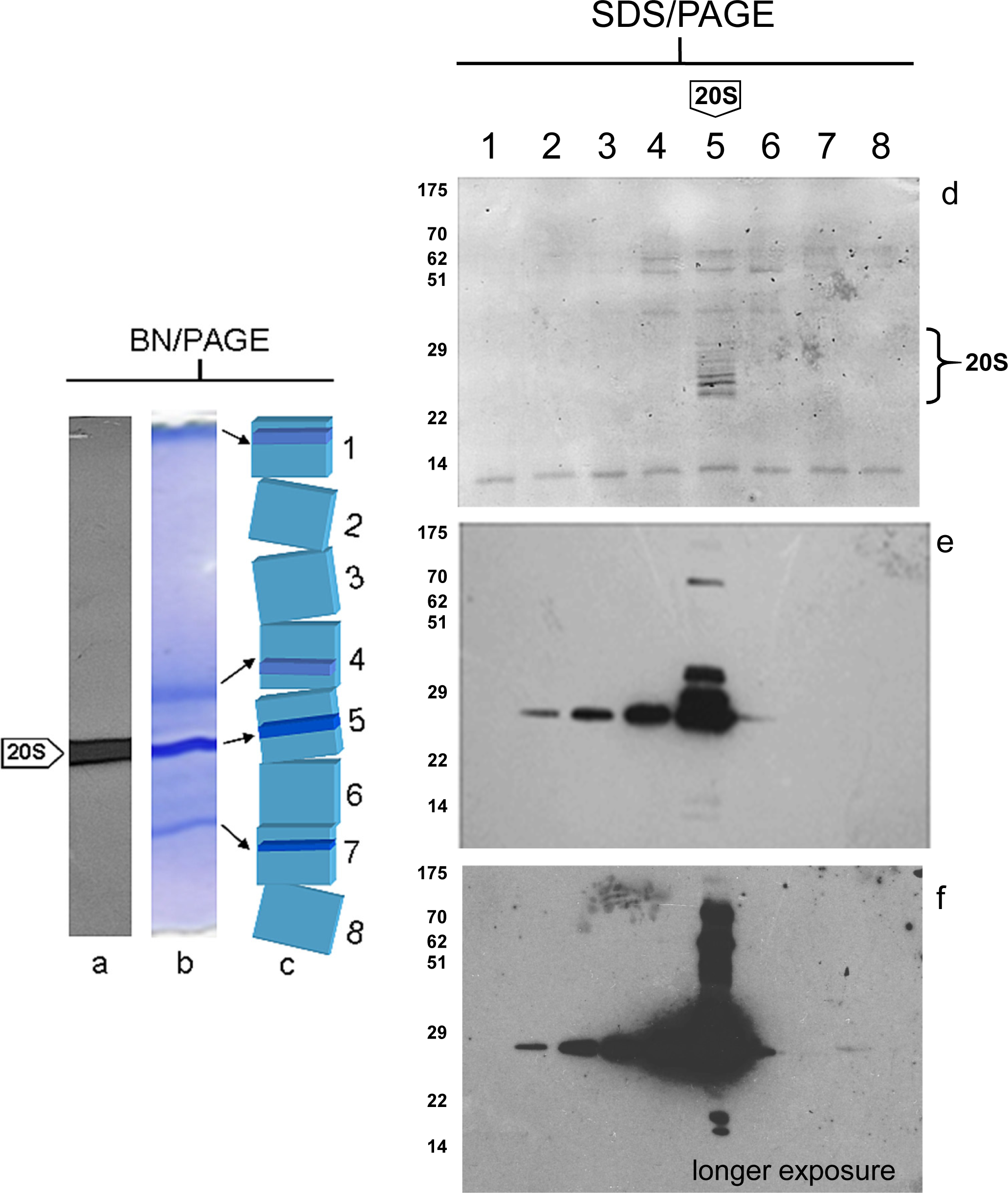
Separation of the different proteasome versions of *A. thaliana* cells by BN/PAGE and analysis by western blot. Proteasome enriched fraction (**P3**) was separated by BN/PAGE (**b**). In-gel denatured proteins were transferred to a nitrocellulose membrane to identify 20S (CP) subunits with an anti-20S antibody (**a**). To detect higher molecular mass proteasomes, a complete BN/PAGE gel was cut into eight fractions (1 to 8, **c**). Proteins contained in each fraction were electroeluted/concentrated to be analyzed independently by SDS/PAGE and western blot using an anti-20S antibody. Transferred proteins were stained with Ponceau S (**d**) before 20S (CP) detection by western blot (**e** and **f**).

### 2.5 SDS/PAGE

We analyzed the protein profile of each electroeluted sample by SDS/PAGE according to the protocol of Laemmli, 1970. Samples in sample buffer were loaded onto 4% stacking gels and resolved in 12% polyacrylamide-SDS gels. Runs were performed using the Tris/glycine/SDS running buffer (see previous section) at 200 V for 1h at 5 °C in a Mini-PROTEAN system (Bio-Rad). Gels were stained in a solution of Coomassie Brilliant Blue R-250 (0.1%), acetic acid (40%) and ethanol (40%). For silver staining of proteins on BN/PAGE, we used the improved method of Blum et al., 1986. [41]. We employed Pink pre-stained ladder, 15-175 kDa (Cat. MWP02, Nippon Genetics) as molecular weight markers.

### 2.6 Western blot and slot blot analysis

Proteins separated by BN/PAGE or SDS/PAGE were electrophoretically transferred to nitrocellulose membranes (0.45 μM HATF, Millipore) using transfer buffer (25 mM Tris, 192 mM glycine, 20% isopropanol) in a Mini-PROTEAN (Bio-Rad) transfer system at 360 mA for 1h at 6 °C. Proteins on BN/PAGE were denatured previous to their blotting by incubating the gel in a solution of 20mM Tris-HCl buffer (pH 7.4), 3% SDS for 10 min with agitation followed by heating in a microwave oven (medium level for 1 min). After an additional incubation for 10 min at room temperature, proteins were transferred as indicated. Proteins on membranes were fixed (25% isopropanol, 10% acetic acid) for 1h with agitation at room temperature. After distilled water wash, proteins were visualized with Ponceau S solution (0.2% Ponceau S in 5% acetic acid). For western blot assay, membranes were blocked 1h at 25 °C with 5% non-fat milk in TBS-T buffer (20mM Tris-HCl pH7.4, 150mM NaCl, 0.05% Tween-20) and incubated with primary or secondary antibodies in the solution of non-fat milk in TBS-T buffer. All intermediate washes were done with TBS-T. The following primary antibodies were used at the same 1:10 000 dilutions for 2 h at 25 °C: mouse-anti-proteasome 20S alpha+beta (Cat. ab22673, Abcam), rabbit-anti-proteasome 26S S2 (Rpn1) (Cat. ab98865, Abcam), rabbit-anti-proteasome Rpn6 (S9) (Cat. PW8370, Enzo), mouse-anti-Rpt2 (Cat. ab21882, Abcam), rabbit-anti-19S S5A/Rpn10 (Cat. ab56851, Abcam) and rabbit-anti-Ubiquitin antibody (Cat. sc-9133, Santa Cruz Biotechnology, Inc.). As secondary antibodies, we employed HRP-goat-anti-mouse IgG (H+L) (Cat. 62-6520, Zymed) or HRP-goat-anti-rabbit IgG (H+L) (Cat. 65-6120, Zymed) both at 1:10 000 dilutions for 1 h at 25 °C. Western blots were developed with Super Signal West Femto (Cat. 34095, Thermo Scientific) and exposed to X-Ray films (Cat. 6040331, Kodak). Total carbonyl (Johansson et al., 2004) and ubiquitin conjugates (Tang et al., 2014) contents of heat stress and control samples, were estimated by slot blot analysis. For carbonyl content estimation, protein from total cell lysates (section 2.2) was precipitated with methanol/chloroform and resuspended in 1X Laemmli sample buffer. Five milligrams of each sample (determined by the method of Lowry, 1951 [44]) were derivatized with 2,4-dinitrophenylhydrazine (DNPH) and loaded on each well of the slot blot manifold (Cat. PR 648, Hoefer). Oxidatively modified proteins on nitrocellulose filters were determined by an immunochemical protocol (OxyBlot Protein Oxidation detection kit, from Chemicon International). Same slot blot technique was followed to determine ubiquitin conjugates, but samples were not derivatized. Quantification of western and slot blots was made by densitometry of the autoradiograms using NIH ImageJ 1.48 software. All data were standardized for growth in control conditions (wild type = 1).

### 2.7 Protein quantification by a modified Lowry method

We determined protein content in those samples that contained β-ME, by the Lowry method according to the modification of Makkar *et al.,* 1980. Briefly, an aliquot of each sample was vacuum dried to eliminate β-ME (from Laemmli sample buffer), which interferes with the protein determination. Samples were resuspended with 0.1N NaOH for its protein quantification by the classic Lowry method with Folin-Ciocalteau reagent calibrated with crystalline bovine serum albumin.

### 2.8 Mass spectrometry

For the mass spectrometric analysis, BN/PAGE slices were distained and chemically modified prior to mass spectrometry analysis. After reduction (dithiothreitol) an alkylation (iodoacetamide) samples were digested in-gel with trypsin (Promega, Madison, WI, USA). Resultant peptides were desalted with Zip Tips C18 (Millipore-Billerica, MA, USA) and applied to a LC-MS system (Liquid Chromatography-Mass Spectrometry) composed by a nanoflow pump (EASY-nLC II, Thermo-Fisher Co. San Jose, CA) and a LTQ-Orbitrap Velos (Thermo-Fisher Co., San Jose, CA) mass spectrometer with a nano-electrospray ionization (ESI) source. The mass spectrometer was calibrated with a Calmix solution containing N-butylamine, caffeine, Met-Arg-Phe-Ala (MRFA), and Ultramark 1621. For LC, a 10%–80% gradient of solution B (water/acetonitrile 0.1% formic acid) was used during 120 min through a home-made capillary column (0.75 µm in diameter × 10 cm in length; RP-C18) with a flux of 300 nL/min. Collision-Induced Dissociation (CID) and High-energy Collision Dissociation (HCD) methods were used for peptide fragmentation, selecting only 2+, 3+ and 4+ charged ions. Single charged ions and those above 5+, as well as ions with undefined charges, were not considered. For data acquisition, a positive ion mode was set. Capture and performance of fragmentation data were done according to the total ion scanning and predetermined charge with 3.0 (m/z) isolation width, a collision energy of 35 arbitrary units, an activation Q of 0.250, an activation time of 10 milliseconds and a maximum injection time of 10 milliseconds per micro-scanning. The automatic capture of data was done using ion dynamic exclusion: (i) exclusion list of 400 ions; (ii) pre-exclusion time of 30 s; and (iii) exclusion time of 300 s. Data were searched against an available *A. thaliana* NCBI databases using Discoverer 1.4 software (Thermo-Fisher Co., San Jose, CA, USA).

### 2.9 Cell viability test

To determine cell viability, samples were collected from the stressed cell cultures at 30 min, 1, 2 and 3 h and from unexposed cells at 0 h and 3 h. For all cultures viability was also quantified after 3 h recovery at 25 ^°^ C. 100 μL of cell culture was mixed with one volume of 0.4 % trypan blue (Sigma-Aldrich) and were incubated for 3 min at 25 ^°^ C. Viable (unstained) and dead (stained) cells were counted in a Neubauer chamber under a light microscope. Cell viability was considered as the percentage of unstained cells out of the total of cells observed.

## 3. Results

### 3.1 Proteasome isolation

The isolation of proteasomes by a protocol of differential centrifugation has been published (Shibatani et al., 2006). This technique has proven effective in capturing all the possible proteasome versions present in a reticulocyte model. We adapted this protocol to isolate the different proteasome versions in suspension cells cultures of *A. thaliana*. The analysis by SDS/PAGE and western blot of representative aliquots (pellets and supernatants) collected along the isolation protocol revealed that the final pellet was effectively enriched in proteasomes (P3, Fig 1b). An enrichment factor (P3/Sn1) of 22.5 X was estimated by film densitometry of a western blot using an anti-20S proteasome.

### 3.2 Separation of different proteasome versions of *A. thaliana* cells by BN/PAGE, concentration by electroelution and protein blot analysis

Since potentially different proteasome versions were contained in the crude P3 fraction from *A. thaliana* cells (Fig. 1b), we loaded this sample directly onto the stacking well of a BN/PAGE for their separation, transfer and detection by western blot with an anti-20S antibody. A Coomassie-stained lateral strip from a BN/PAGE, revealed a profile of four major bands of a molecular mass (estimated by the method described by Wittig *et al. 2006*) of 560, 750, 850 and 2600 kDa (Fig. 2b). Western blot from transferred native gels shown that only one of these bands with a molecular mass of ∼ 750 kDa, had a strong reactivity with the anti-20S antibody (Fig. 2a). The migration of this band was consistent with the native molecular mass of the independent 20S (CP) by BN/PAGE (Shibatani et al., 2006; Camacho-Carvajal et al., 2004). Nevertheless, our western blot analysis failed at showing proteasome complexes of a molecular mass higher than the 20S (CP) (Fig. 2a). One possibility to explain this result might be the low abundance of superior proteasomes versions or the limitation of our detection system. To circumvent this problem, we cut an entire mini BN/PAGE (8 × 6 cm) into eight horizontal fractions that were independently electroeluted and analyzed (Fig. 2c). One of the major advantages of electroelution in the system we used, in addition to its quantitative sample recovery (Ohhashi et al., 1991; Sui et al., 1996), is that it concentrates the contained proteins. For the protocol here described, we estimated a concentration factor of 35X. The components of disassembled and electroeluted proteasomes in every gel piece (section 2.4) were resolved by SDS/PAGE and analyzed by western blot using the anti-20S antibody. Figure 2d shows a representative profile of the proteins recovered from a whole BN/PAGE and immobilized on a nitrocellulose filter. Ponceau S stain evidenced an abundant set of bands between 20 and 30 kDa (lane 5, Fig. 2d) electroeluted from the fraction 5 that contained the band originally recognized by the anti-20S antibody when an intact native gel was transferred (Fig. 2a). The analysis of an equivalent gel fraction by mass spectrometry (Supplemental Table 1) indicated that this band corresponded to the independent 20S (CP). Additional evidence that proteins in fraction 5 corresponded to the 20S (CP) subunits, was given by the anti-20S antibody (Fig. 2e and f). A longer film exposure to the same western blot membrane (Fig. 2f) produced a heavy smear in the 40 to 100 kDa interval. Western blot analysis on a 20S (CP) purified from an equivalent enriched proteasome fraction (P3, Fig. 1) by ion exchange chromatography and size exclusion fractionation (Supplemental Fig. 1), suggests that the subunits of the 20S are heavily ubiquitinated even though the proteasome fraction was purified from *A. thaliana* cells kept under the optimal culture conditions (Fig. 2e and f). By Ponceau S staining (or Coomassie staining on an equivalent gel) the characteristic 20S (CP) set of bands were hard to observe in those fractions that potentially contained proteasomes with a molecular mass higher than the independent 20S (CP) (lanes 1 to 4, Fig. 2d). Nevertheless, the anti-20S antibody tracked proteasomes up to lanes 2, 3 and 4 (Fig. 2e and f) that correspond to putative proteasome complexes of a nominal average molecular mass (estimated by BN/PAGE) of approximately 1600, 1100 and 850 kDa, respectively. The proteasomes profile described in Fig. 2 was considered as the basal for *A. thaliana* suspension cells under optimum culture conditions as defined in this paper, where the predominant population of proteasomes was constituted by the independent 20S (CP) and the abundance of “heavier” proteasomes gradually decreased toward the top of the BN/PAGE (Fig. 2e and f).

### 3.3 High molecular mass proteasomes in heat-stressed cells

Same general methodology (proteasome isolation, separation of different proteasome versions by BN/PAGE, electroelution/concentration and western blot) was applied to *A. thaliana* cells exposed to heat stress to detect possible alterations in the basal proteasomes arrangement observed in control cells (Fig. 2). First, we needed to establish if the content of total 20S proteasomes changed at 37 °C exposition. Densitometry analysis of the western blot films showed a small increment in the anti-20S antibody signal from 11 to 17% between unexposed cells and those at 37 °C and less of an 8% among stressed cells (Supplementary Fig. 2a to c). Since 20S content among samples were considered equivalent, for comparative purposes the BN/PAGE of proteasome-enriched fraction (P3) prepared from all culture cells were loaded with the same protein content (Fig. 3). After 30 min of heat treatment, a major difference was detected on fraction 1 (Lane 1, Fig. 3c). A faint signal produced by a proteasome population of a presumed molecular mass of ∼ 2 600 kDa (based on its BN/PAGE mobility) was detected. The signal from the proteasomes versions contained in fractions 2 to 4 was equivalent to the unexposed cells (lanes 2 to 4 Fig. 3a and c). We also noticed the presence of signal bands between 50 to 100 kDa (lane 5 Fig. 3c) that probably correspond to ubiquitinated subunits of the 20S proteasome (Supplementary Fig. 1b). Parallel determinations were also carried out to establish the global levels of protein ubiquitination and carbonylation of total cell lysates (Fig. 4). Both parameters have been broadly used as markers of cellular stress (Lledías et al., 1999; Taylor et al., 2002; Bollineni et al., 2014). At 30 min of heat treatment we detected the removal of 10% of the original total ubiquitin conjugates content by slot blot analysis (Fig. 4a) and a slight ubiquitination signal clearance in the western blot image (lane 2, Supplementary Fig. 2d). The level of oxidatively modified proteins on the other hand, showed no changes at this time (Fig. 4b). The trypan blue cell viability assay showed no difference between the control and heat stressed cells. Trypan blue exclusion in stressed cells was equivalent to control culture even after 3 h recovery at 25 ^°^ C. A striking difference in the western blot proteasomes profile was detected at 1 and 2 h after the temperature increase, where an important enrichment of the higher order proteasome configurations was observed (lanes 1 to 4, Fig. 3d and e). In addition, a “new” anti-20S signal was detected in the fraction 6 from these heat-stressed cells cultures (lane 6 Fig. 3d and e) originated from a native protein complex of approximately 640 kDa. This molecular mass was significantly smaller than a functional 20S (CP). We speculated that in this fraction, because of the reactivity with the anti-20S (lane 6, Fig. 3d to f), the estimated native molecular mass by BN/PAGE and the 20S peptides (α and β) obtained by mass spectrometry (not shown), 20S assembly intermediate complexes known as half-proteasomes (13 - 16S) could be localized (Schmidtke et al., 1997; Lehmann et al., 2002). The 50 - 100 kDa smear of the 20S (CP) subunits was equivalent at both times (lane 5, Fig. 3d and e) suggesting a strong 20S (CP) subunits ubiquitination. The increment in high molecular mass proteasome populations correlated with the clearance of 60% (at 1h) and 75% (at 2h) of the basal ubiquitinated proteins levels (Fig. 4a and Supplementary Fig. 2d). The total amount of protein carbonyls was practically unaltered at these times (Fig. 4b). Cell viability test showed no changes with respect to control cultures when determined immediately after 37 ^°^ C treatment or the recovery for 3 h at 25 ^°^ C was allowed. The western blot analysis of cell suspension cultures at 3 h under heat stress, showed the higher enrichment of all the proteasome versions for all the times sampled, half-proteasomes included (Fig. 3 f). In addition, a noteworthy feature of this 3 h profile was the presence of 20S-immunoreactive bands between 60 and 70 kDa in all fractions (bracket in Fig. 3f, lanes 1 to 8). These bands are presumably produced by the ubiquitinated subunits of the 20S (CP) (Supplemental Fig. 1) that assembled the half-proteasomes, the free 20S CP and the higher molecular mass proteasome versions promoted by heat stress. There were also clear differences in the slot blot determination of total ubiquitin conjugates and carbonyl contents, a two-fold and a nine-fold increase respectively in comparison with the previous sampled hour (Fig. 4a and b). No changes in cell viability were observed relative to control cultures at 3h heating or after the recovery period.

**Fig. 3.**
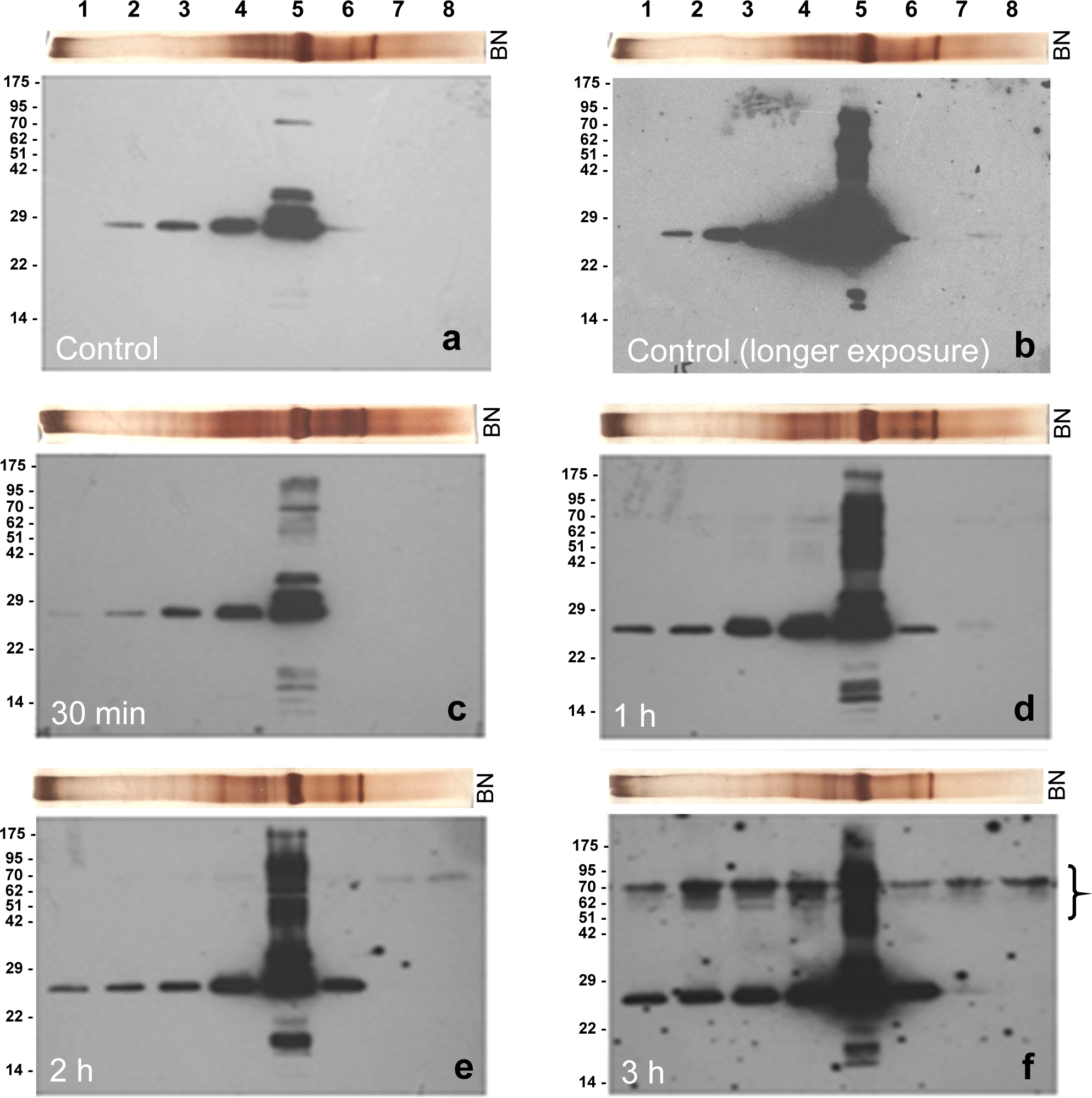
Western blot proteasomes profiles of *A. thaliana* cells under heat stress. Proteasome enriched fractions (**P3**) from cells cultures at 37°C were separated by BN/PAGE. Eight fractions obtained from each BN/PAGE gel (lanes 1 to 8) were individually electroeluted/concentrated, precipitated and analyzed by western blot using an anti-20S antibody. Panel **a** correspond to unexposed cells, **c** to **e** show the profiles of the cells recovered at 30 min, 1, 2 and 3 h, respectively. Except for **b**, all films were exposed the same time to the chemiluminescent developing reaction. Above to each film image, we positioned a silver stained BN/PAGE lane (**BN**) to show the actual protein content and profile of each P3 sample from where the eight fractions were obtained. The most abundant protein in all BN/PAGE, electroeluted from fraction 5 (lane 5, a to f) was the independent 20S (CP). The signals from fractions 1 to 4 (**a** to **f**) were produced by proteasome versions with a molecular mass higher than the 20S (CP).

**Fig. 4.**
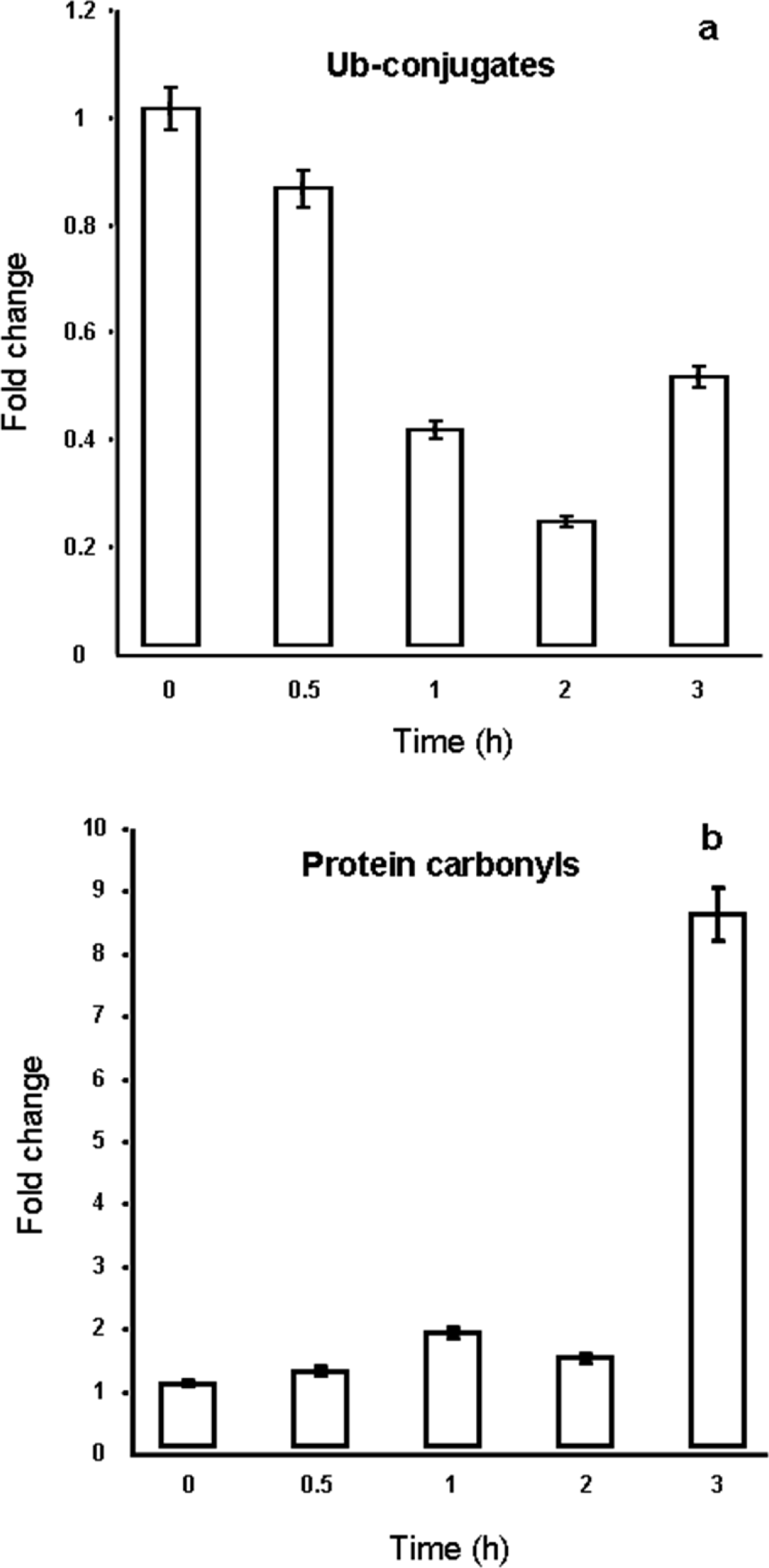
Ubiquitin conjugates and protein carbonyls of total lysates from *A. thaliana* cell cultures exposed to heat stress. 5 mg of total protein extract were loaded into slot blot wells and immobilized in nitrocellulose membranes for the determination of the total amount of ubiquitin conjugates present in cells incubated at 37 °C for 0.5, 1, 2 and 3 h with a monoclonal anti-Ubiquitin antibody (**a**). Same protocol was followed to determine the content of oxidatively modified proteins, except samples were derivatized with DNPH before their membrane immobilization (**b**). Detection was made with an anti-DNPH antibody. Densitometry films values were normalized to 1, considering the ubiquitin conjugates or total carbonyl contents from an unexposed control (0 h). Media and standard deviations of three independent experiments are represented in the graphs.

### 3.3 19S regulatory particle subunits are part of the high molecular mass proteasomes

The modular nature of proteasomes and their association/dissociation dynamics directed by environmental clues has been reported (Bajorek et al., 2003; Grune et al., 2011; Livnat-Levanon et al., 2014). High molecular mass proteasomes resolved by BN/PAGE were result from the assembly of different regulators on the distal surface of one or both the 20S α rings (Shibatani et al., 2006). To determine if the high molecular mass proteasomes we observed contained 19S regulatory particle subunits, we probed the eight fractions obtained from a control culture BN/PAGE with antibodies against Rpn1 (19S base subunit), Rpt2 (19S base subunit), Rpn10 (19S lid subunit that keeps together base and lid) and Rpn6 (lid subunit which holds together 20S and 19S complexes). Fraction 5 that contained exclusively the independent 20S (CP) (lane 5, figure 5b and c) showed minimum or null reactivity against all the anti-19S regulatory subunits antibodies (lane 5, Fig. 5d to g). A strong signal with the immediate higher molecular mass proteasome complex was obtained with the Rpn10 and Rpn1 antibodies (Lane 4, Fig. 5d and e). Rpn10 signal showed a stepwise decrease toward the upper region of the BN/PAGE (Lanes 1 to 5, Fig. 5d) while the Rpn1 signal kept constant in three consecutive fractions and showed a decrease up to fraction 1 (Lanes 2 to 4, Fig. 5e). The use of the Rpt2 and Rpn6 antibodies shown a very different pattern, a stepwise increase toward those fractions obtained from the top of the BN/PAGE. If we consider that, all the subunits are found in the context of their respective 20S or 19S complex (Livneh et al, 2016), our results suggest that some of the detected complexes probably represent 26S maturation/assembly intermediates. Based on the BN/PAGE mobility of the complex and the relative reactivity of the antibodies in each fraction, Rpn10 and Rpn1 were associated with the 20S CP (figure 5d and e, lanes 3 and 4) before association of Rpt2 and Rpn6 (Fig. 5f and g, lanes 3 and 4) as has been described (Hendil et al., 2009). In this context BN/PAGE fractions (Lanes 3 and 4, Figure 5) could be considered early steps in the way to consolidate the higher molecular mass proteasome complexes 20S/19S and 19S/20S/19S probably contained in fractions 2 and 1 respectively (Lane 2 and 1, Fig. 5).

**Fig. 5.**
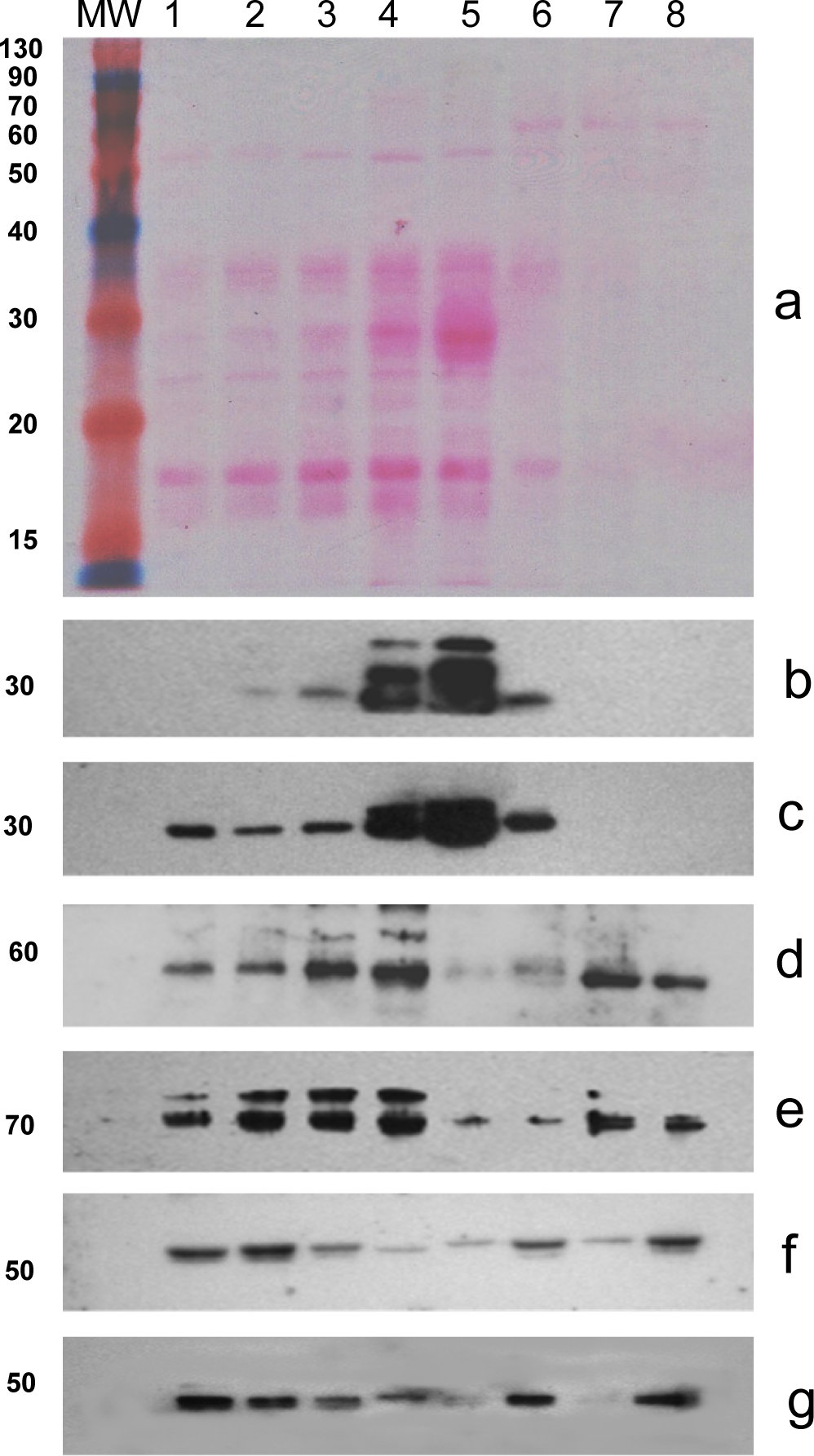
19S regulatory particle subunits were present in high molecular mass proteasomes. Denatured BN/PAGE fractions (1-8) from control *A thaliana* cells were transferred to nitrocellulose, stained with Ponceau red (**a**) and probed with anti-20S (**b** and **c**), anti-Rpn 10 (**d**), anti-Rpn 1 (**e**), anti-Rpt 2 (**f**) and anti-Rpn 6 (**g**) antibodies. In contrast with panel (**b)**, the anti-20S profile in **c**, was obtained with five times higher protein loading.

## 4. Discussion

Different proteasomes versions have been shown by BN/PAGE analysis of samples from a human embryonic cell line (HEK293) (Camacho-Carvajal et al., 2004) and rabbit reticulocytes [34] (Shibatani et al., 2006). The separation of the different proteasomes versions by BN/PAGE revealed that stimulation with γ-interferon or MG132 (a proteasome inhibitor) directed a dynamic process of recruitment and exchange of proteasome regulatory complexes (Shibatani et al., 2006). We reasoned that the published protocols of proteasomes isolation and electrophoretic analysis could be applicable to *A. thaliana* suspension cells, and as a starting point, to detect whether different proteasome versions are present in plants. We also would be able to analyze the proteasome populations changes in response to an insult such as temperature increment. In our hands, the published protocol for proteasome isolation (Shibatani et al., 2006) with some modifications was effective for *A. thaliana* cells in suspension cultures (Fig. 1b). In our lab, the same technique was useful for the isolation of proteasomes (and their BN/PAGE analysis) from *A. thaliana* two-week seedlings, mature spinach and maize leaves, different succulent plants leaves and from yeast, mouse liver, zebra fish and human erythrocytes (Rivas and Lledías, unpublished). When the enriched proteasome fraction (P3) of *A. thaliana* cells was separated by BN/PAGE and transferred to membranes for western blot detection with an anti-20S antibody, only one band was detected (Fig. 2a). Higher molecular mass complexes were not observed at this point. We discarded the possibility that in our adaptation of the isolation protocol, any of the buffers or additives employed or even the sample freezing, promoted the disassembly of complex proteasomes. The same enrichment protocol has been successfully used as a previous step to purify, by ionic exchange chromatography and size exclusion fractionation, the 26S proteasomes from *A. thaliana* suspension culture cells. An alternative possibility was that the abundance of higher order proteasomes was relatively scarce in this cell type and were beyond the limit of detection. The electroelution protocol was effective for concentrating the proteasomes in all BN/PAGE fractions and facilitate their visualization by western blot (Fig. 2 and 3). Four proteasome populations were revealed in suspension cells grown at optimum culture conditions (25 °C). The most abundant proteasome version corresponded to the independent 20S (CP) (Fig. 2 e and f). The identity of the 20S (CP) was verified by mass spectrometric analysis (Supplementary Table 1). We discarded the possibility that “heavier” proteasomes versions detected in unexposed or in heat stressed cells (Fig. 3) were product of an artifact caused by anomalous or not optimum electrophoretic separation of independent 20S (CP) particles, since fractions 1 to 6 (Fig. 3) electroeluted under native conditions and independently re-separated on fresh blue native gels, were detected at exactly the same original fractions. Even though samples were frozen at −70 °C after their electroelution, high molecular mass complexes, the independent 20S (CP) and lower complexes preserved their original electrophoretic mobility. We observed that in cells, even at optimum growth conditions, the subunits of the 20S (CP) were probably ubiquitinated (Supplemental Fig. 1 and smear in lane 5 Fig. 3b). The posttranscriptional modification of the 20S (CP) subunits by ubiquitination have been shown by proteomic techniques in *A. thaliana* (Book et al., 2010). High temperatures by themselves caused intracellular protein denaturation and substrates ubiquitination (Lepock et al., 1988; Pinto et al., 1991). In *A. thaliana* suspension cell cultures, a moderate heat stress was detected when the temperature was raised at 37 °C and the production of reactive oxygen species (ROS) was enhanced (Volkov et al., 2006) that in turn may promote protein carbonylation. A classical marker of cellular oxidative stress is the increase of carbonyls in total protein samples (Levine et al., 1990; Wong et al., 2010; Bollineni et al., 2014). Oxidation partially denatures protein and hydrophobic patches exposure initializes the intricate action of the ubiquitin-proteasome system (UPS) (Pacifici et al., 1993; Murata et al., 2001). Protein ubiquitin conjugates are considered an early and sensitive cell stress marker (Shang and Taylor 2011). In our experiments, the levels of ubiquitin conjugates and oxidatively modified proteins suggest two phases in the *A. thaliana* cell response to the heat increment. During the first phase (30 min to 2 h at 37 °C) 75 % the basal ubiquitin conjugates were removed (Fig. 4a and Supplementary Fig. 2d) while the total protein carbonyl level remained relatively unaltered (Fig. 4b). These results indicate that an oxidative stress was not produced because the antioxidant machinery and the modified protein elimination mechanisms (ub-conjugates degradation among them) were effective. The western blot proteasome profiles (Fig. 3a to e) suggest that heat increment promoted the assembly of proteasome versions of a molecular mass higher than the free 20S (CP). In reticulocytes the differences in molecular mass of proteasomes complexes detected by BN/PAGE have been attributed to the interaction of the 20S (CP) with the 19S particle to constitute the 26S proteasome (Shibatani et al., 2006; Camacho-Carvajal et al., 2004) which in turn is responsible of ubiquitin conjugates elimination (Smalle and Vierstra, 2004; Voges et al., 1999) while the still independent 20S (CP) degrades oxidatively modified proteins (Ferrington et al., 2001; Grune et al., 1997). In *A. thaliana,* the abundance of these two proteasome entities is highly interrelated during cell growth and stress tolerance (Kurepa et al., 2009). If the degradation of protein ubiquitin conjugates was limited (by 26S synthesis impairment) the degradation of oxidized proteins by the independent 20S (CP) was favored (Kurepa et al., 2008). We consider that the opposite phenomenon as we observe during the first phase is also plausible. During the second phase (3 h at 37 °C) both stress markers increased, doubled for ubiquitin conjugates and a nine-fold increase was detected for protein oxidation (Fig. 4a and b). These results are indicative that cellular antioxidant and damaged protein removal capacities were surpassed, and an oxidative stress episode was initiated (Sies, 1997). The western blot profile showed the higher enrichment for all the proteasomes versions observed (Fig. 3 f). The stress markers levels and the western blot results combined, suggest that high molecular mass proteasomes still removed ubiquitin conjugates but with a relative diminished efficiency (Fig. 4a); however, oxidized proteins greatly increased (Fig.4b). We hypothesize that high temperature for 3 h, forced the cell machinery to assembly high molecular mass proteasomes while decreasing the total amount of independent 20S (CP), oxidatively modified proteins increased consequently. The 26S proteasome has been shown inefficient at degrading oxidatively modified proteins (Davies, 2001) and in addition, was inhibited by oxidative stress (Reinheckel et al., 1998). This observation probably explains why despite “heavier” proteasomes were enriched at this time (Fig. 3f) a decrease of 50% in their capability of ubiquitinated proteins removal was detected (Fig. 4a). An additional explanation for the impaired oxidized protein elimination is the sensitivity of the trypsin and caspase-like 20S (CP) catalytic activities to oxidation (Demasi et al., 2003). A heavy signal, attributable to 20S subunits ubiquitination was observed for all the BN/PAGE fractions from suspension cells heat stressed for 3h (Fig. 3f and Supplemental Fig. 1). Under abiotic stress, the activity of specific *A. thaliana* ubiquitin ligases (Pub22 and Pub23) increased, which destabilized the 19S regulator by ubiquitination of its subunits (Cho et al., 2015). In human embryonic cells (HEK293) the ubiquitination of the subunit Rpn13 (a ubiquitinated proteins receptors located in the regulatory particle 19S) decreased its substrate binding capacities under proteotoxic stress conditions leading to ubiquitin conjugates build up (Besche et al., 2014). Inhibition of the 26S activity by substrate overload is plausible (Kurepa et al., 2009). All these processes are probably happening in cells in suspension cultures throughout 3 h of heat exposition.

Severe heat stress causes an increase in ubiquitinated proteins (Ferguson et al., 1994) and the accumulation of protein carbonyls has been reported as a product of heat exposition in plants that produced an excess in ROS production (Hasanuzzaman et al., 2013). In our experiments, the elimination of ubiquitin conjugates and the avoidance of the accumulation of protein oxidation products during the first 2 h of heat exposition suggest that *A. thaliana* suspension cell cultures adapted the proteasome degradative machinery (and additional cellular mechanisms) to tolerate the temperature increment, a tolerance that seemed limited to the third hour where the cell stress markers increased (Fig 4a and b). Despite this fact, cell viability values remain unaltered for 3 h, a parameter that indicated that at least for this period, the cell capacity to cope with heat stress was not entirely compromised.

The films in Fig. 3 (a, c to f) were exposed the same time to the chemiluminescence developing reaction to allow a direct comparison among the different proteasome populations present at different times under stress. The contrast between the proteasome complement of control cells with those of the stressed cultures (Fig. 3), even when a longer film exposure was practiced for control cells (Fig. 3b), strongly suggests that heat stress promoted the assembly of proteasome versions with a molecular mass higher than the independent 20S (CP). Our western blot analysis of total lysates showed that there was not a significant net increase in the total content of proteasomes 20S (CP) subunits among the heat-treated cells (Supplemental Fig. 2). However, the profiles obtained by the separation of different proteasomes populations by BN/PAGE (Fig. 3) and 20S ubiquitination (Supplementary Fig. 1), suggest that synthesis, assembly and probably degradation of proteasomes subunits were promoted under heat stress. During this very dynamic process, proteasomes subunits could be synthesized and assembled as half-proteasomes through their activation as mature 20S (CP). Eukaryotic half-proteasomes are assembly complexes constituted by a seven-membered α ring and several β subunits proproteins (Schmidtke et al., 1997; Lehmann et al., 2002 and dedicated chaperones (Le Tallec et al., 2007). In line with our observation, mammalian half-proteasomes were localized just above the band of mature 20S (CP) by native gel electrophoresis (Schmidtke et al., 1997). Once 20S (CP) are completed, they are available as platforms to assembly higher molecular mass proteasomes versions, or they can remain independent, in close dependence with the protein turn over needs imposed by the environment. The western blot analysis of the BN/PAGE fractions 1 to 8 from control cells, showed that the 19S regulatory particle subunits Rpn10, Rpn1, Rpt2 and Rpn6 are associated to “supra” 20S proteasome assemblies (Fig. 5). The use of antibodies against the 19S regulatory particle subunits Rpn10, Rpn1 and Rpt2 produced a faint signal associated with the independent 20S and the putative semi-proteasome (Fig. 5, lanes 5 and 6 respectively). Signals from non-20S associated forms were detected in fractions 7 and 8 (Fig. 5d to g). We propose that some of the observed heat-enriched proteasome versions, corresponded to the 26S assembly and other alternative proteasome versions with ub-conjugates elimination capacities, while the isolated 20S (CP) kept oxidatively modified proteins at basal levels. We cannot discard the possibility that even other high molecular mass proteasome forms were also involved in the ubiquitin conjugates and oxidized proteins elimination. Experimental evidenced have shown that during an oxidative stress event in yeast, Ecm29 increased its association with the 19S regulatory complex promoting 26S disassembly. Since independent 20S (CP) population increased oxidized modified proteins could be preferentially degraded (Wang et al., 2010). In murine fibroblasts, proteasomes with an array PA28-20S-PA28 induced during oxidative stress, degraded carbonylated proteins with a higher efficiency (Pickering et al., 2010).

The presence in the cell of *ad hoc* proteasomes could offer better possibilities for successfully coping with unfavorable growth conditions. Additional and higher resolution analyses are needed to identify the specific protein components in each fraction in the Ub-proteasome pathway context, nevertheless we consider that our approach of proteasome isolation by centrifugation, separation of discrete proteasomes populations based in their molecular mass differences by BN/PAGE, and the concentration of the samples by electroelution shown that an increment in culture temperature directed the assembly of “supra” 20S proteasome complexes. Our protocol (combined with mass spectrometry and western blot) could be considered a useful tool to characterize the regulators and the additional interacting proteins that contribute to the proteasomes function and dynamics.

## Contribution

Daniel Aristizábal: performed the experiments and analyzed the data. Viridiana Rivas: performed some of the experiments. Fernando Lledías: conceived and designed the experiments, performed some of the experiments and wrote the manuscript. Gladys Cassab: wrote the manuscript.

## Acknowledgments

We are thankful to P. Guzmán and L. Aguilar (CINVESTAV, Irapuato) for providing the *A. thaliana* suspension cell cultures. We also thank Unidad de Proteómica, Instituto de Biotecnología, Universidad Nacional Autónoma de México for the mass spectrometry analysis.

## Ethical approval

This article does not contain any studies with human participants or animals performed by any of the authors.

## Abbreviations

BN/PAGE: Blue Native Polyacrylamide Gel Electrophoresis
SDS/PAGE: SDS Polyacrylamide Gel Electrophoresis

## REFERENCES

Bajorek M, Finley D, Glickman MH. 2003. Proteasome disassembly and downregulation is correlated with viability during stationary phase. Current Biology 13, 1140–1144

Beck F, Unverdorben P, Bohn S, Schweitzer A, Pfeifer G, Sakata E, Nickell S, Plitzko JM, Villa E, Baumeister W, Forster F. 2012. Near-atomic resolution structural model of the yeast 26S proteasome. Proceedings of the National Academy of Sciences 109, 14870–14875

Besche HC, Sha Z, Kukushkin NV, Peth A, Hock EM, Woong, Gygi S, Gutierrez JA, Liao H, Dick L, Goldberg AL. 2014. Autoubiquitination of the 26S Proteasome on Rpn13 Regulates Breakdown of Ubiquitin Conjugates. EMBO Journal 33, 1159– 1176

Blickwedehl J, Olejniczak S, Cummings R, Sarvaiya N, Mantilla A, Chanan-Khan A, Pandita TK, Schmidt M, Thompson CB, Bangia N. 2012. The proteasome activator PA200 regulates tumor cell responsiveness to glutamine and resistance to ionizing radiation. Molecular Cancer Research 10, 937–944

Blum H, Beier H, Gross HJ. 1986. Improved silver staining of plant proteins, RNA and DNA in polyacrylamide gels. Electrophoresis 8, 93–99

Bohn S, Beck F, Sakata E, Walzthoeni T, Beck M, Aebersold R, Förster F, Baumeister W, Nickell S. 2010. Structure of the 26S proteasome from Schizosaccharomyces pombe at subnanometer resolution. Proceedings of National Academy of Sciences 107, 20992–20997

Bollineni RC, Hoffmann R, Fedorova M. 2014. Proteome-wide profiling of carbonylated proteins and carbonylation sites in HeLa cells under mild oxidative stress conditions. Free Radical Biology and Medicine 68, 186–95

Book AJ, Gladman NP, Lee SS, Scalf M, Smith LM, Vierstra RD. 2010. Affinity purification of the Arabidopsis 26S proteasome reveals a diverse array of plant proteolytic complexes Journal of Biological Chemistry 285, 25554–25569

Camacho-Carvajal MM, Wollscheid B, Aebersold R, Steimle V, Schamel WW. 2004. Two-dimensional Blue native/SDS gel electrophoresis of multi-protein complexes from whole cellular lysates: a proteomics approach. Molecular and Cellular Proteomics 3, 176–182.

Cho SK, Bae H, Ryu MY, Wook Yang S, Kim WT. 2015. PUB22 and PUB23 U-BOX E3 ligases directly ubiquitinate RPN6, a 26S proteasome lid subunit, for subsequent degradation in Arabidopsis thaliana Biochemical and Biophysical Research Communication 464, 994-999

Couttas T, Raftery M, Erce M, Wilkins M. 2011. Monitoring cytoplasmic protein complexes with blue native gel electrophoresis and stable isotope labelling with amino acids in cell culture: analysis of changes in the 20S proteasome. Electrophoresis 32, 1819–1823.

Davies KJ. 2001. Degradation of oxidized proteins by the 20S proteasome. Biochimie 83, 301–10.

Demasi M, Silva GM, Netto LE. 2003. 20S proteasome from Saccharomyces cerevisiae is responsive to redox modifications and is S-glutathionylated. Journal of Biological Chemistry 278, 679–685

Ferguson IB, Lurie S, Bowen JH. 1994. Protein synthesis and breakdown during heat shock of cultured pear (Pyrus communis) cells. Plant Physiology 104, 1429–1437

Fort P, Andrey V, Kajava AV, Delsuc F, Coux O. 2015. Evolution of Proteasome Regulators in Eukaryotes. Genome Biology and Evolution 7, 1363–1379.

Ferrington DA, Sun H, Murray KK, Costa J, Williams TD, Bigelow DJ, Squier TC. 2001. Selective degradation of oxidized calmodulin by the 20 S proteasome. Journal of Biological Chemistry 276, 937–943

Glickman MH, Ciechanover A. 2002. The ubiquitin-proteasome proteolytic pathway: Destruction for the sake of construction. Physiological Reviews 82, 373–428

Groll, M. Bajorek M, Köhler Moroder L, Rubin DM Huber R, Glickman MH, Finley D. 2000. A gated channel into the proteasome core particle. Nature Structural Biology 7, 1062–1067

Grune T, Reinheckel T, Davies KJ. (1997) Degradation of oxidized proteins in mammalian cells. FASEB Journal 11, 526–534

Grune T, Catalgol B, Licht A, Ermak G, Pickering AM, Ngo JK, Davies KJ. 2011. HSP70 mediates dissociation and reassociation of the 26S proteasome during adaptation to oxidative stress. Free Radicals Biology and Medicine 51, 1355–64

Hasanuzzaman M, Nahar K, Alam MM, Roychowdhury R, Masayuki Fujita M. 2013. Physiological, biochemical, and molecular mechanisms of heat stress tolerance in plants. International Journal of Molecular Sciences 14, 9643–9684

Hashemi A, Gharechahi J, Nematzadeh G, Shekari F, Hosseini SA, Salekdeh GH 2016. Two-dimensional blue native/SDS-PAGE analysis of whole cell lysate protein complexes of rice in response to salt stress. Journal of Plant Physiology 200, 90–101

Hendil KB, Kriegenburg F, Tanaka K, Murata S, Lauridsen AB, Johnsen AH, Hartmann-Petersen R 2009. The 20S proteasome as an assembly platform for the19S regulatory complex. Journal of Molecular Biology 394, 320–328

Hernebring M, Fredriksson Å, Liljevald M, Cvijovic M, Norman K, Wiseman J, Semb H, Nyström T. 2013. Removal of damaged proteins during ES cell fate specification requires the proteasome activator PA28. Scientific Reports 3, 1–6

Hershko A, Ciechanover A. 1998. The ubiquitin system. Annual Review of Biochemistry 67, 425–79

Johansson E, Olsson O, Nystrom T. 2004. Progression and specificity of protein oxidation in the life cycle of Arabidopsis thaliana. Journal of Biological Chemistry 279, 22204–22208.

Kish-Trier E, Hill CP. 2013. Structural biology of the proteasome. Annual Review of Biophysics 42, 29–49

Kurepa J, Toh-e A, Smalle JA. 2008. 26S proteasome regulatory particle mutants have increased oxidative stress tolerance. Plant Journal 53, 102–114

Kurepa J, Wang S, Li Y, Smalle J. 2009. Proteasome regulation, plant growth and stress tolerance. Plant Signaling and Behavior 4, 924–927

Laemmli UK. 1970. Cleavage of Structural Proteins during the Assembly of the Head of Bacteriophage. Nature 227, 680–685

Lasserre JP, Beyne E, Pyndiah S, Lapaillerie D, Claverol S, Bonneu M. 2006. A complexomic study of Escherichia coli using two-dimensional blue native/SDS polyacrylamide gel electrophoresis. Electrophoresis 27, 3306–21

Le Tallec B, Barrault MB, Courbeyrette R, Guérois R, Marsolier-Kergoat MC, Peyroche A. 2007. 20S proteasome assembly is orchestrated by two distinct pairs of chaperones in yeast and in mammals. Molecular Cell 27, 660–674

Lehmann A, Janek K, Braun B, Kloetzel PM, Enenkel C. 2002. 20S proteasomes are imported as precursor complexes into the nucleus of yeast. Journal of Molecular Biology 317, 401–413

Lehmann A, Niewienda A, Jechow K, Janek K, Enenkel C. 2010. Ecm29 fulfills quality control functions in proteasome assembly. Molecular Cell 38, 879-888

Lepock JR, Frey HE, Rodahl AM, Kruuv J. 1988. Thermal analysis of CHL V79 cells using differential scanning calorimetry: implications for hyperthermic cell killing and the heat shock response. Journal of Cellular Physiology 137, 14–24

Levine RL, Garland D, Oliver CN, Amici A, Climent I, Lenz AG, Ahn BW, Shaltiel S, Stadtman E R. 1990. Determination of carbonyl content in oxidatively modified proteins Methods in Enzymology 186, 464–478

Livneh I, Cohen-Kaplan V, Cohen-Rosenzweig C, Avni N, Ciechanover A. 2016. The life cycle of the 26S proteasome: from birth, through regulation and function, and onto its death. Cell Research 26, 869–885

Livnat-Levanon N, Kevei E, Kleifeld O, Krutauz D, Segref A, Rinaldi T, Erpapazoglou Z, Cohen M, Reis N, Hoppe T. 2014. Reversible 26S proteasome disassembly upon mitochondrial stress. Cell Reports 7, 1371–1380

Lledías F, Rangel P, Hansberg W. 1999. Singlet oxygen is part of a hyperoxidant state generated during spore germination. Free Radical Biology and Medicine 26, 1396–404

Lowry OH, Rosebrough NJ, Farr AL, Randall RJ. 1951. Protein measurement with the Folin phenol reagent. Journal of Biological Chemistry 193, 265–275

Makkar H, Sharma O, Negi S. 1980. Assay of proteins by Lowry’s method in the presence of high concentrations of beta-mercaptoethanol. Analytical Biochemistry 104, 124–116

Manzano C, Abraham Z, López-Torrejón G, Del Pozo JC. 2008. Identification of ubiquitinated proteins in Arabidopsis. Plant Molecular Biology 68, 145-58

Murashige T, and Skoog F. 1962. A revised medium for rapid growth and bioassays with tobacco tissue cultures. Physiologia Plantarum 15, 473–497

Murata S, Minami Y, Minami M, Chiba T, Tanaka K. 2001. CHIP is a chaperone-dependent E3 ligase that ubiquitylates unfolded protein. EMBO Reports 2, 1133–1138

Nussbaum, AK Dick TP, Keilholz W, Schirle M, Stevanović S, Dietz K, Heinemeyer W, Groll M, Wolf DH, Huber R, Rammensee HG, Schild H. 1998. Cleavage motifs of the yeast 20S proteasome β subunits deduced from digests of enolase 1. Proceedings of the National Academy of Sciences 95, 12504–12509

Obin M, Shang F, Gong X, Handelman G, Blumberg J, Taylor A. 1998. Redox regulation of ubiquitin-conjugating enzymes: mechanistic insights using the thiol-specific oxidant diamide. FASEB Journal 12, 561–569

Ohhashi T, Moritani C, Andoh H, Satoh S, Ohmori S, Lottspeich F, Ikeda M. 1991. Preparative high-yield electroelution of proteins after separation by sodium dodecyl sulphate-polyacrylamide gel electrophoresis and its application to the analysis of amino acid sequences and to raise antibodies. Journal of Chromatography 585, 153– 159

Pacifici RE, Kono Y, Davies J. 1993. Hydrophobicity as the signal for selective degradation of hydroxyl radical-modified hemoglobin by the multicatalytic proteinase complex, proteasome. Journal of Biological Chemistry 268, 15405–15411

Pickering AM, Koop AL, Teoh CY, Ermak G, Grune T, Davies KJ. 2010. The immunoproteasome, the 20S proteasome and the PA28αβ proteasome regulator are oxidative-stress-adaptive proteolytic complexes. Biochemical Journal 432, 585–594

Polge C, Jaquinod M, Holzer F, Bourguignon J, Walling L, Brouquisse R. 2009. Evidence for the existence in Arabidopsis thaliana of the proteasome proteolytic pathway activation in response to cadmium. Journal of Biological Chemistry 284, 35412–35424

Peth A, Uchiki T, Goldberg AL. 2010. ATP-dependent steps in the binding of ubiquitin conjugates to the 26S proteasome that commits to degradation. Molecular Cell 40, 671–681

Pinto M, Morange M, Bensaude O. 1991. Denaturation of proteins during heat shock. In vivo recovery of solubility and activity of reporter enzymes. Journal of Biological Chemistry 266, 13941–13946

Reinheckel T, Sitte N, Ullrich O, Kuckelkorn U, Davies KJ, Grune T. 1998. Comparative resistance of the 20S and 26S proteasome to oxidative stress. Biochemical Journal 335, 637–642

Sakata E, Bohn S, Mihalache O, Kiss P, Beck F, Nagy I, Nickell S, Tanaka K, Saeki Y, Forster F, Baumeister W. 2012. Localization of the proteasomal ubiquitin receptors Rpn10 and Rpn13 by electron cryomicroscopy. Proceedings of National Academy of Sciences 109, 1479–1484

Schmidtke G, Schmidt, Kloetzel PM. 1997. Maturation of mammalian 20S proteasome: purification and characterization of 13S and 16S proteasome precursor complexes. Journal of Molecular Biology 268, 95–106

Schweitze A, Aufderheide A, Rudack T, Beck F, Pfeifer G, Plitzko JM, Sakata E, Schulten K, Forster F, Baumeister W. 2016. Structure of the human 26S proteasome at a resolution of 3.9 A. Proceedings of National Academy of Sciences 113, 7816-7821

Sessler N, Krug K, Nordheim A, Mordmuller B, Macek B. 2012. Analysis of the Plasmodium falciparum proteasome using Blue Native PAGE and label-free quantitative mass spectrometry. Amino Acids 43, 1119–1129

Shang F, Taylor A. 1995. Oxidative stress and recovery from oxidative stress are associated with altered ubiquitin-conjugating and proteolytic activities in bovine lens epithelial cells. Biochemical Journal 307, 297–303

Shang F, Taylor A. 2011. Ubiquitin–proteasome pathway and cellular responses to oxidative stress. Free Radical Biology and Medicine 51, 5–16

Shibatani T, Carlson EJ, Larabee F, McCormack AL, Früh K, Skach WR. 2006. Global organization and function of mammalian cytosolic proteasome pools: implications for PA28 and 19S regulatory complexes. Molecular Biology of the Cell 17, 4962–4971

Shibahara T, Kawasaki H, Hirano H. 2002. Identification of the 19S regulatory particle subunits from the rice 26S proteasome. European Journal of Biochemistry 5, 1474–83.

Sies H. 1997. Oxidative stress: oxidants and antioxidants. Experimental Physiology 82, 291–295

Smalle J, Vierstra RD. 2004. The ubiquitin 26S proteasome proteolytic pathway. Annual Review of Plant Physiology 55, 555–590

Sui LM, Hughes W, Hoppe AJ, Petra PH. 1996. Direct evidence for the localization of the steroid-binding site of the plasma sex steroid-binding protein (SBP or SHBG) at the interface between the subunits. Protein Science 5, 2514–2520

Tanaka K, Yoshimura T, Kumatori A, Ichihara A, Ikai A, Nishigai M, Kameyama K, Takagi T. 1988. Proteasomes (multi-protease complexes) as 20S ring-shaped particles in a variety of eukaryotic cells Journal of Biological Chemistry 263, 16209-16217

Tang CH, Leu MY, Shao K, Hwang LY, Chang WB. 2014. Short-term effects of thermal stress on the responses of branchial protein quality control and osmoregulation in a reef-associated fish, Chromis viridis. Zoological Studies 53, 1–9.

Taylor A, Shang FT, Nowell T, Galanty Y, Shiloh Y. 2002. Ubiquitination capabilities in response to neocarzinostatin and H_2_O_2_ stress in cell lines from patients with ataxia-telangiectasia. Oncogene 21, 4363–4373

Thompson AR, Vierstra RD. 2005. Autophagic recycling: lessons from yeast help define the process in plants. Current Opinion in Plant Biology 8, 165–173

Volkov R, Panchuk I, Mullineaux P, Schoffl F. 2006. Heat stress-induced H_2_O_2_ is required for effective expression of heat shock genes in Arabidopsis. Plant Molecular Biology 61, 733–746.

Voges D, Zwickl P, Baumeister, W. 1999. The 26S proteasome: a molecular machine designed for controlled proteolysis. Annual Review of Biochemistry 68, 1015–1068

Wang YJ, Kaiser P, Huang L. 2010. Regulation of the 26S proteasome complex during oxidative stress. Science Signaling 3, 1–10

Wittig I, Braun HP, Schägger H. 2006, Blue native PAGE. Nature Protocols 1, 418– 28

Wittig I, Beckhaus T, Wumaier Z, Karas M, Schägge H. 2010. Mass estimation of native proteins by blue native electrophoresis. Principles and practical hints. Molecular and Cellular Proteomics 9, 2149–2161

Wong CM, Marcocci L, Liu L, Suzuki YJ. 2010. Cell signaling by protein carbonylation and decarbonylation. Antioxidants and Redox Signaling 12, 393–404

Yang P, Fu H, Walker J, Papa CM, Smalle J, Ju YM, Vierstra RD. 2004. Purification of the Arabidopsis 26 S proteasome: biochemical and molecular analyses revealed the presence of multiple isoforms. Journal of Biological Chemistry 279, 6401– 6413

