## Supplemental Fig 1 for "Heat stress reveals high molecular mass proteasomes in *Arabidopsis thaliana* suspension cells cultures"

Departamento de Biología Molecular de Plantas

Instituto de Biotecnología, Universidad Nacional Autónoma de México

Av. Universidad 2001, Col. Chamilpa, Cuernavaca, Mor., 62250, México.

*Corresponding Author

**Supplementary Fig. 1. 20S (CP) subunits ubiquitination.** 100 μg of protein (proteasome enriched fraction, P3, Fig. 1) from cells kept under optimal growth conditions were fractionated on a 1 mL UnoQ1 (Bio-Rad) anion exchange column using a 0-1M HCl gradient in Tris 20 mM (pH 7.4). Fractions (1 mL) with activity for Suc-LLVY-AMC chymotrypsin substrate plus 0.03% SDS were pooled, dialyzed, concentrated (Amicon centricon YM30) and resolved on a Superose 6, 10/300GL (Pharmacia) in Tris 20 mM (pH7.4). Four of the central activity "peak" fractions (lanes 1 to 4) were precipitated with methanol/chloroform and prepared for their analysis by SDS/PAGE and western blot using an anti-20S antibody (**a**) or a monoclonal anti-ubiquitin antibody (**b**).

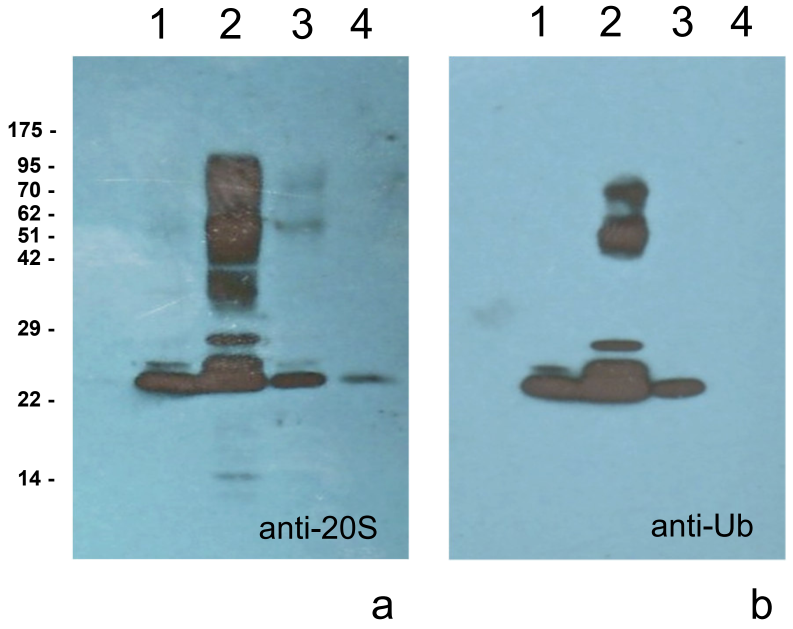


**Supplementary Fig. 1**
