## Supplemental Fig 2 for "Heat stress reveals high molecular mass proteasomes in *Arabidopsis thaliana* suspension cells cultures"

Departamento de Biología Molecular de Plantas

Instituto de Biotecnología, Universidad Nacional Autónoma de México

Av. Universidad 2001, Col. Chamilpa, Cuernavaca, Mor., 62250, México.

*Corresponding Author

**Supplementary Fig. 2. Total content of 20S proteasome subunits and ubiquitin conjugates in lysates of *A. thaliana* cells exposed to heat stress.** 20 μg of protein was loaded per lane for its SDS/PAGE separation, (coomassie stained) (**a**), protein transferred to nitrocellulose membrane (Ponceau S stained) (**b**), 20S (CP) subunits detection by an anti-20S antibody (**c**). The numbers at the bottom of each lane correspond to the film densitometry analysis. Ub conjugates using an anti-Ub monoclonal antibody (**d**). Lane 1, unexposed cells. Lanes 2 to 5, cells at 37 $^{o}$C for 30min, 1, 2 and 3 h respectively.

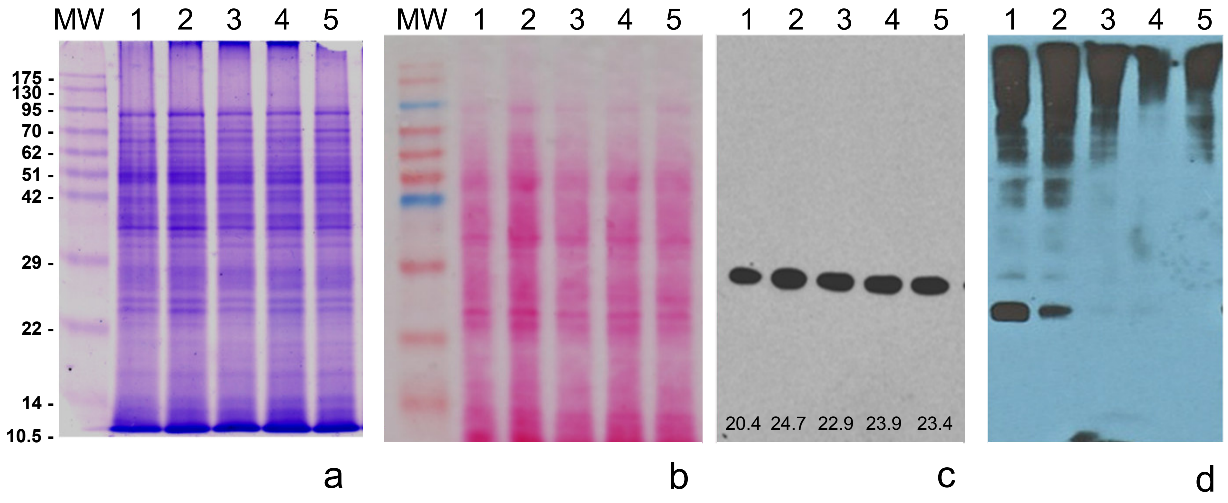


**Supplementary Fig. 2**
