## Supplemental Table 1 for "Heat stress reveals high molecular mass proteasomes in *Arabidopsis thaliana* suspension cells cultures"

**Supplementary Table 1.** Proteins identified by mass spectrometry analysis following BN/PAGE and electroelution/concentration of fraction 5 from *A. thaliana* cells

| Rank | Uniq Pep | Acc # | Lledias5.dta/Lledias5 Num Unique | % Cov | Best Disc Score | Best Expect Val | Protein MW | Species | Protein Name |
|---|---|---|---|---|---|---|---|---|---|
| 1 | | 15220961 | 10 | 48.9 | 5.28 | 3.5e-9 | 25947.5 | ARABIDOPSIS THALIANA | PAE1; endopeptidase/ peptidase/ threonine-type endopeptidase |
| 1-1 | 2 | 15231824 | 10 | 48.9 | 5.28 | 3.5e-9 | 25977.5 | ARABIDOPSIS THALIANA | PAE2; endopeptidase/ peptidase/ threonine-type endopeptidase |
| 2 | | 15222152 | 9 | 32.1 | 7.17 | 1.1e-12 | 29667.8 | ARABIDOPSIS THALIANA | PBE1; endopeptidase/ peptidase/ threonine-type endopeptidase |
| 3 | | 15220151 | 9 | 37.9 | 4.65 | 5.3e-8 | 30410.2 | ARABIDOPSIS THALIANA | PAF2; endopeptidase/ peptidase/ threonine-type endopeptidase |
| 4 | | 15233268 | 7 | 36.0 | 5.61 | 8.7e-10 | 27475.5 | ARABIDOPSIS THALIANA | PAC1; endopeptidase/ peptidase/ threonine-type endopeptidase |
| 5 | | 15225839 | 7 | 32.9 | 3.23 | 2.8e-6 | 27377.6 | ARABIDOPSIS THALIANA | PAG1; endopeptidase/ peptidase/ threonine-type endopeptidase |
| 6 | | 15232965 | 5 | 26.0 | 4.31 | 2.2e-7 | 24644.3 | ARABIDOPSIS THALIANA | PBF1; peptidase/ threonine-type endopeptidase |
| 7 | | 15223537 | 5 | 28.5 | 4.35 | 1.9e-7 | 27651.6 | ARABIDOPSIS THALIANA | PBG1; peptidase/ threonine-type endopeptidase |
| 8 | | 15219257 | 5 | 24.7 | 3.53 | 6.2e-6 | 25701.5 | ARABIDOPSIS THALIANA | PAB1 (PROTEASOME SUBUNIT PAB1); endopeptidase/ peptidase/ threonine-type endopeptidase |
| 9 | | 15230435 | 5 | 19.2 | 5.53 | 1.3e-9 | 27337.4 | ARABIDOPSIS THALIANA | PAD1 (20s proteasome alpha subunit pad1); endopeptidase/ peptidase/ threonine-type endopeptidase |
| 9-1 | 1 | 15239271 | 5 | 19.2 | 3.25 | 6.3e-6 | 27324.3 | ARABIDOPSIS THALIANA | PAD2 (PROTEASOME ALPHA SUBUNIT D 2); endopeptidase/ peptidase/ threonine-type endopeptidase |
| 9-2 | 1 | 2511580 | 5 | 20.4 | 3.25 | 6.3e-6 | 25845.6 | ARABIDOPSIS THALIANA | multicatalytic endopeptidase |
| 10 | | 15235889 | 5 | 21.5 | 4.86 | 2.1e-8 | 25151.6 | ARABIDOPSIS THALIANA | PBA1; endopeptidase/ peptidase/ threonine-type endopeptidase |
| 11 | | 15238554 | 4 | 17.9 | 4.35 | 1.9e-7 | 27294.3 | ARABIDOPSIS THALIANA | PAA1 (PROTEASOME ALPHA SUBUNIT A 1); endopeptidase/ peptidase/ threonine-type endopeptidase |
| 11-1 | 1 | 15224993 | 3 | 11.8 | 1.91 | 4.5e-4 | 27350.3 | ARABIDOPSIS THALIANA | PAA2 (20S PROTEASOME SUBUNIT PAA2); endopeptidase/ peptidase/ threonine-type endopeptidase |
| 11-2 | 1 | 79322198 | 3 | 13.2 | 1.91 | 4.5e-4 | 24456.2 | ARABIDOPSIS THALIANA | PAA2 (20S PROTEASOME SUBUNIT PAA2); endopeptidase/ peptidase/ threonine-type endopeptidase |
| 12 | | 15228805 | 5 | 25.0 | 3.50 | 3.2e-6 | 22540.9 | ARABIDOPSIS THALIANA | PBD1 (20S PROTEASOME BETA SUBUNIT D1); peptidase/ threonine-type endopeptidase |
| 12-1 | 1 | 15233580 | 4 | 18.1 | 1.95 | 4.4e-4 | 21984.3 | ARABIDOPSIS THALIANA | PBD2 (20S PROTEASOME BETA SUBUNIT 2); peptidase/ threonine-type endopeptidase |
| 13 | | 18395025 | 3 | 20.1 | 4.17 | 4.1e-7 | 22798.4 | ARABIDOPSIS THALIANA | PBC1 (PROTEASOME BETA SUBUNIT C1); peptidase/ threonine-type endopeptidase |
| 14 | | 15237451 | 3 | 9.9 | 2.32 | 3.8e-5 | 29617.3 | ARABIDOPSIS THALIANA | PBB2; endopeptidase/ peptidase/ threonine-type endopeptidase |
| 14-1 | 1 | 18405364 | 3 | 9.9 | 2.32 | 3.8e-5 | 29555.2 | ARABIDOPSIS THALIANA | PBB1; endopeptidase/ peptidase/ threonine-type endopeptidase |
| 14-2 | 1 | 2511578 | 3 | 9.9 | 2.32 | 3.8e-5 | 29524.1 | ARABIDOPSIS THALIANA | multicatalytic endopeptidase |
| 14-3 | 1 | 227204261 | 2 | 7.5 | 2.32 | 3.8e-5 | 27435.7 | ARABIDOPSIS THALIANA | AT3G27430 |
| 14-4 | 1 | 30688785 | 2 | 7.1 | 2.32 | 3.8e-5 | 28812.1 | ARABIDOPSIS THALIANA | PBB1; endopeptidase/ peptidase/ threonine-type endopeptidase |
| 15 | | 14532710 | 1 | 1.1 | 1.98 | 3.5e-5 | 108882.6 | ARABIDOPSIS THALIANA | unknown protein |
